# Survey of rice (*Oryza sativa* L.) production ecosystems in northern Ghana confirms low risk of exposure to potential toxic elements from local grain consumption

**DOI:** 10.1101/2023.04.08.536104

**Authors:** Eureka E. A. Adomako, Kow Aboagye-Ghunney, Prince Owusu

## Abstract

Expanding local rice production to meet consumer demand is a priority action under the Government of Ghana’s Planting for Food and Jobs initiative. While studies on yield-enhancing interventions including seed improvement and fertilizer management abound, fewer studies focus on food safety issues such as the potential toxic element status of the production ecosystems. This study was, therefore, conducted to bridge the knowledge gap. Chemical analyses were conducted on water, soil and rice grain samples from rainfed upland, rainfed lowland and irrigated lowland rice ecosystems in the Northern and Upper East regions of the country. Statistical analysis of the data showed that soil and rice grain arsenic concentrations were significantly higher (P < 0.001) in the Upper East region. In the Northern region, mean cadmium concentration in rice grains from the irrigated lowland fields (0.023 ± 0.003 mg/kg) was significantly higher than in grains from the rainfed fields. All recorded concentrations of rice grain arsenic, cadmium and lead were, however, within permissible limits, indicating a low risk of dietary exposure. The observed differences in concentrations within and between regions suggest that soil texture and other geogenic factors could influence the potential toxic element status of the rice production ecosystems. Regular monitoring is, therefore, recommended to maintain the safety of Ghana’s locally produced rice for human consumption.

## 1. Introduction

Rice (*Oryza sativa* L.) is an important staple for more than half of the world’s population (GRiSP, 2013) and is considered as a priority crop under the Planting for Food and Jobs (PFJ) initiative launched by the Government of Ghana in 2017 (Pauw, 2022). It is grown in all the agro-ecological zones and administrative regions of Ghana, with regions in the northern part of the country – notably, Northern, Upper East and Upper West – accounting for half of local production (MoFA-IFPRI, 2020). Despite a steady growth of about 11% per annum between 2008 and 2019, local rice production in Ghana continues to fall about 50% short of consumer demand (MoFA-IFPRI, 2020) due to an increasing shift in dietary preferences from the indigenous staples (mainly cassava, yam and maize) to rice. This has been attributed to urbanization, the ease of cooking rice compared to other local dishes, and changes in consumer habit as people become wealthier (Al-Hassan et al., 2008; Anabila, 2015; MoFA-IFPRI, 2020). Per capita consumption of milled rice in Ghana was estimated to have increased from 13.3 kg/annum in 1990 to 32.0 kg/annum in 2015 (SRID-MoFA, 2017) and was projected to reach 45 kg/annum by 2025 (Van Oort et al., 2015). SRID-MoFA (2020), however, reported an increase in per capita consumption of milled rice from 40.4 kg/annum in 2016 to 51.5 kg/annum in 2020; thus, exceeding the earlier projection by Van Oort et al. (2015). According to Kushitor (2021), rice has replaced maize as the primary cereal in Ghana. Studies have also shown a higher preference for imported rice than locally produced rice (Anabila, 2015; Graham-Acquaah et al., 2020; Ragasa et al., 2020). Factors that determine consumers’ choice include taste, appearance (good-looking grains), aroma, absence of foreign materials and packaging (Adomako et al., 2011; Ehiakpor et al., 2017).

To address concerns regarding foreign exchange imbalances due to high dependence on rice imports and Ghana’s vulnerability to price volatility in the international rice market, the National Rice Development Strategy of 2009 (reviewed in 2015) and the government’s PFJ initiative both aim at expanding local rice production to meet consumer demand (MoFA-IFPRI, 2020; Onumah et al., 2022; Pauw, 2022; Ragasa and Chapoto, 2017). National food and agriculture policy interventions directed at meeting the ambitious targets for local rice output have focused on facilitating access to farm inputs, mainly improved seed and fertilizers (Pauw, 2022; Tanko et al., 2019; Zakaria et al., 2021). Rice research in Ghana is, therefore, skewed towards seed improvement interventions aimed at enhancing yield and pest/disease resistance (Abebrese et al., 2019; Fordjour et al., 2020; Tawiah et al., 2022). Efforts have also been made to better understand the effects of different fertilizer/nutrient management regimes (Fuseini and Benjamin, 2022; Sarkodee-Addo et al., 2021) and the socio-economic factors that influence farmers’ adoption of improved rice varieties and other productivity enhancing technologies (Azumah et al., 2022; Tanko, 2022; Tanko and Ismaila, 2021; Quaye et al., 2022). In contrast, fewer studies have been conducted on the status of potential toxic elements (PTEs), e.g. arsenic (As), cadmium (Cd) and lead (Pb), in Ghana’s rice production ecosystems. Such studies are, however, necessary since consumers can only derive the full benefits of the nutritional value of this important food staple when it is safe for consumption.

In other parts of the world - particularly in Asia - several studies have been conducted in rice production systems, all of which point to environmental regulation as a key determinant of potential toxic element (PTE) concentrations in rice grain (Adomako et al., 2009; Ali et al., 2020; Lu et al., 2009; Williams et al., 2006). To date, the study by Adomako et al. (2010) is the largest field survey of PTE concentrations in Ghanaian rice production ecosystems. It did not, however, include rice fields in northern Ghana due to logistical constraints. This study was, therefore, conducted to expand the scope to include rice fields in northern Ghana, build on the existing data and validate earlier findings regarding the safety of Ghana’s locally produced rice for human consumption.

## 2. Method

### 2.1. Study sites

Field surveys were conducted in the Tolon District of the Northern Region and the Kassena-Nankana East Municipality of the Upper East Region of Ghana. A total of eighteen (18) study sites representing three types of rice production ecosystems in northern Ghana - rainfed upland (RU), rainfed lowland (RL) and irrigated lowland (IL) - were surveyed. The selected study sites were all cultivated with AGRA rice, an aromatic variety widely grown in northern Ghana.

In the Northern Region, three RU and three RL sites were selected from research fields belonging to the Council for Scientific and Industrial Research – Savanna Agricultural Research Institute (CSIR-SARI) at Nyamkpala and three IL sites were selected from fields around the Golinga Dam. In the Upper East region, three IL sites were selected from fields around the Tono Dam, along with three RU and three RL sites belonging to local farmers in Navrongo. Communication with the local farmers and CSIR-SARI scientists revealed that fertilizers were applied in all the study sites to enhance plant growth.

### 2.2 Sample collection

At each sampling site, soil samples were collected using a 5-cm diameter soil auger to core beside the base of mature rice plants to a depth of 20 cm. Matured rice grains (110-140 days after sowing) were also harvested from the same locations as the soil samples. The grain samples were placed in brown paper envelopes while soil samples were put into resealable polyethylene bags. All the envelopes and bags were sealed and appropriately labelled to ensure correct matching of soil and grain samples from the same location.

At the IL study sites located around the Golinga and Tono Dams, water samples were collected from irrigation canals supplying the rice fields into 500-ml plastic bottles that had been pre-washed with detergent, soaked and washed in 10% nitric acid (HNO_3_) for 24 hours, rinsed thoroughly with double distilled water and oven dried overnight (Eaton et al., 2005). In the Northern region, additional water samples were collected from Bontanga (a rice irrigation site close to Golinga); and in the Upper East Region, reference samples were obtained from a borehole dug for domestic purposes at Navrongo. Prior to collection of the water samples, on-site rinsing of the plastic bottles was done three times with the water to be sampled. For total recoverable metal analysis, each sample was acidified with 5 ml of HNO_3_ to a pH below 2.0 according to USEPA Method 3005 (Edgell, 1989). The collected water samples were sealed and appropriately labelled to indicate the sampling location and sample number. They were immediately stored in an ice chest at 4°C to protect against the effects of light and extreme heat that could alter the water chemistry by enhancing microbial activity. On transfer to the laboratory, the water samples were kept in a refrigerator at 4°C for 48 hours pending analysis.

### 2.3 Sample preparation and analysis

Soil samples were air-dried at room temperature for 5 – 7 days, then disaggregated with a porcelain mortar and pestle, and sieved to 2 mm prior to analysis. Soil physicochemical properties – namely pH, electrical conductivity (EC) and particle size distribution – were all determined using standard protocols (Van Reeuwijk, 2002). Rice grain samples were manually de-husked and oven-dried at 90°C for 48 hours in a brown envelope until constant dry weight. They were then milled into powdery form in a blender and sieved to 1 mm. Water samples were taken out of the freezer 24 hours prior to analysis and allowed to thaw at room temperature.

#### 2.3.1 Digestion of water samples

For determination of total As, Cd and Pb concentrations in the water samples, 40 ml of each sample was measured into a 250-ml digestion flask and 15 ml of nitric acid (HNO_3_) were added before heating on a block digester (Behrotest^®^ K8) at 45°C for 1-2 hours. Each sample digest was then filtered on cooling with a Whatman No. 42 filter paper into a 100-ml volumetric flask and topped up to the mark with de-ionized water.

#### 2.3.2 Digestion of soil samples

For total element detection in the soils, 1.0 g of each sample was measured into a digestion tube to which 10 ml of ternary mixture – 500 ml of HNO_3_, 50 ml of concentrated sulphuric acid (H_2_SO_4_) and 20 ml of perchloric acid (HClO_4_) – were added and then placed on a hot plate at 95 °C under a fume hood for 30 minutes. After cooling, the soil digests were filtered with a Whatman No. 42 filter paper into 100-ml volumetric flasks and topped up to the mark with deionized water.

#### 2.3.3 Digestion of rice samples

For determination of total rice grain As, Cd and Pb concentrations, 0.5 g of each milled and sieved sample was weighed into a 100 ml conical flask and 5 ml of (H_2_SO_4_) was added and left to react overnight. Subsequently, the samples were placed on a hot sand bath at 95 °C and, after about 20 minutes, hydrogen peroxide (H_2_O_2_) was added dropwise until the samples were clear in colour indicating complete digestion. On cooling, the rice grain digests were also filtered into 100-ml volumetric flasks and topped up to the mark as already described.

#### 2.3.4 Determination of elemental concentrations

Total elemental concentrations in all the samples were measured using the PerkinElmer PINAAcle 900T Atomic Absorption Spectrophotometer (AAS). Cd and Pb concentrations were determined using flame atomic absorption spectrophotometry (FAAS) while As was determined using the flow injection analysis - atomic absorption spectrometry (FIA-AAS) (hydride generation) technique. Air-acetylene gas was used as the source of fuel for Cd and Pb while argon gas was used for As. Table 1 shows the instrument analytical conditions for the respective elements.

**Table 1.** Instrument analytical conditions for investigated elements.

| Spectrometer parameter | As | Cd | Pb |
| --- | --- | --- | --- |
| Wavelength (nm) | 193.7 | 228.8 | 283.3 |
| Slit width (nm) | 0.5 | 0.7 | 0.7 |
| Lamp current (mA) | 10 | 8 | 10 |
| Fuel | Argon | Acetylene | Acetylene |
| Support | Air | Air | Air |

#### 2.3.5 Quality control

For quality assurance, standards prepared from stock solutions of each element, as well as blanks, were used to calibrate the AAS prior to analysis. This was repeated after every set of ten (10) sample measurements to ensure precision and accuracy of the analytical results. As part of the quality control protocol, duplicate samples of a standard reference material (SRM) of moist clay – WEPAL (Wageningen Evaluating Programs for Analytical Laboratories) International Soil-Analytical Exchange Reference material (ISE Sample 999) – were included in every batch of soil sample digestion and analysis in order to guarantee the accuracy of the data. The certified and measured concentrations of the respective elements in the SRM are presented in Table 2.

**Table 2.** Certified and measured concentrations of As, Cd and Pb in SRM (ISE Sample 999)

| Analyte | Certified value (mg/kg) | Measured value (mg/kg) | % Recoveries |
| --- | --- | --- | --- |
| As | $0.772 \pm 0.344$ | $0.852 \pm 0.223$ | 95.3 |
| Cd | $0.040 \pm 0.020$ | $0.048 \pm 0.031$ | 99.6 |
| Pb | $4.710 \pm 1.157$ | $4.907 \pm 0.235$ | 98.7 |

### 2.4 Data Analysis

The Gen-Stat statistical package (version 12) was used for all statistical analyses. Comparison of means for the different rice production ecosystems was by one-way analysis of variance (ANOVA) at a significance level of 5% (*P* ≤ 0.05). Comparison of data obtained for the two regions was by a simple *t* – test, also at a 5% significance level. Post Hoc tests were conducted for data with *P* ≤ 0.05 using Fisher’s Least Significant Difference.

## 3. Results

### 3.1 Soil Physicochemical properties

Data on soil pH, electrical conductivity (EC) and particle size distribution for the various rice production ecosystems in the Northern and Upper East region are presented in Table 3. The lowest mean pH of soils from the Northern Region was recorded at the IL field sites (5.4 ± 0.3) while the highest was at the RL field sites (6.1 ± 0.7). Similarly, soils from the IL field sites in the Upper East region had the lowest mean pH of 5.1 ± 0.1. Statistical analysis of the data showed no significant difference in soil pH across the different production ecosystems (Table 3). As with soil pH, no significant difference was observed in mean EC values across the three rice production ecosystems in both the Northern and Upper East regions (Table 3).

**Table 3.** Physicochemical properties of soil from different rice production ecosystems.

| Soil Property | Region | Rice Production system |  |  | *P*-value |
| --- | --- | --- | --- | --- | --- |
|  |  | Rainfed Upland (n = 3) | Rainfed Lowland (n = 3) | Irrigated Lowland (n = 3) |  |
| pH | Northern | 5.6 ± 0.3 | 6.1 ± 0.7 | 5.4 ± 0.3 | 0.248 |
|  | Upper East | 5.3 ± 0.2 | 5.2 ± 0.2 | 5.1 ± 0.1 | 0.387 |
| EC (µScm-1) | Northern | 55.0 ± 7.4 | 76.7 ± 41.8 | 85.3 ± 42.3 | 0.572 |
|  | Upper East | 127.8 ± 73.3 | 55.8 ± 29.2 | 61.8 ± 15.6 | 0.190 |
| % Sand | Northern | 18.0 ± 0.6<sup>a</sup> | 16.3 ± 2.5<sup>a</sup> | 58.7 ± 8.4<sup>b</sup> | < 0.001 |
|  | Upper East | 23.7 ± 3.1 | 28 ± 5.3 | 26 ± 1.0 | 0.391 |
| % Silt | Northern | 73.0 ± 0.6<sup>a</sup> | 74.7 ± 2.9<sup>a</sup> | 33.0 ± 8.7<sup>b</sup> | < 0.001 |
|  | Upper East | 47 ± 3.0 | 43.3 ± 3.1 | 44.3 ± 2.9 | 0.361 |
| % Clay | Northern | 9.0 ± 1.2 | 9.0 ± 1.7 | 8.3 ± 0.6 | 0.761 |
|  | Upper East | 29.3 ± 2.5 | 28.7 ± 2.3 | 29.7 ± 2.1 | 0.867 |
| Texture | Northern | Silt Loam | Silt Loam | Sandy Loam | - |
|  | Upper East | Clay Loam | Clay Loam | Clay Loam | - |

Means in the same row with the same superscript are not statistically different at $P \leq 0.05$

With respect to particle size distribution, no significant difference was observed in soils from the Upper East Region, all of which had a clay loam texture. In the Northern region, however, a significant difference (*P* < 0.001) was observed in the sand and silt composition of soils from the different production ecosystems. The soil texture at the RU and RL field sites was silt loam, while at the IL field sites it was sandy loam (Table 3).

### 3.2 Elemental concentrations in water samples

Mean concentration of Pb was below detection limit in all the water samples, except for those from Tono which had a mean Pb concentration of 0.011 mg/l (Table 4). Mean As concentration ranged from 0.004 ± 0.002 mg/l in water from the Navrongo borehole to 0.006 ± 0.003 mg/l in water samples collected from the Tono irrigation site, but differences were not statistically significant (Table 4). Although Cd concentrations in water samples from the Upper East region were below detection, detectable levels were recorded in samples from the Bontanga and Golinga IL fields in the Northern region (Table 4).

**Table 4.** Elemental concentrations in water from rice irrigation channels and borehole.

| Sampling Site | Region | As<br>(mg/l) | Cd<br>(mg/l) | Pb<br>(mg/l) |
| --- | --- | --- | --- | --- |
| Golinga<br>(n=3) | Northern | $0.005 \pm 0.001$ | $0.008 \pm 0.003$ | BD |
| Bontanga<br>(n=3) | Northern | $0.005 \pm 0.001$ | $0.009 \pm 0.004$ | BD |
| Tono<br>(n=3) | Upper East | $0.006 \pm 0.003$ | BD | $0.011 \pm 0.019$ |
| Navrongo<br>Borehole<br>(n=2) | Upper East | $0.004 \pm 0.002$ | BD | BD |
| *P*-value |  | 0.070 | - | - |

BD: Below detection

### 3.3 Elemental concentrations in rice field soils

The mean concentrations of As, Cd and Pb in soils sampled from the various rice production ecosystems in the Northern and Upper East regions are presented in Table 5. For both the Northern and Upper East regions, one-way ANOVA of the soil As data showed that differences in the mean concentrations across the three production ecosystems were not statistically significant (Table 5). In the Upper East region, a mean soil As concentration of 0.053 mg/kg was recorded in all the production ecosystems. This was about 3 times higher than the highest mean soil As content (0.016 ± 0.012 mg/kg) recorded in the Northern region IL fields (Table 5). A simple *t*-test found the differences between the two regions with regard to soil As concentration to be significant at *P* < 0.001.

**Table 5.** Elemental concentrations in soil from different rice production ecosystems.

| Element | Region* | Soil PTE concentration in mg/kg |  |  | *P*-value |
| --- | --- | --- | --- | --- | --- |
|  |  | Rainfed Upland | Rainfed Lowland | Irrigated Lowland |  |
| As | Northern | $0.003 \pm 0.001$ | $0.006 \pm 0.004$ | $0.016 \pm 0.012$ | 0.458 |
| | Upper East | $0.053 \pm 0.003$ | $0.053 \pm 0.005$ | $0.053 \pm 0.003$ | 0.985 |
| Cd | Northern | $0.019 \pm 0.010$ | $0.014 \pm 0.003$ | $0.042 \pm 0.019$ | 0.309 |
|  | Upper East | BD | BD | BD | - |
| Pb | Northern | $0.394 \pm 0.156$ | $0.843 \pm 0.087$ | $0.600 \pm 0.268$ | 0.282 |
| | Upper East | $0.193 \pm 0.159$ | $0.614 \pm 0.236$ | $0.432 \pm 0.138$ | 0.270 |

BD: Below detection

While Cd concentration was below detection limit in all the soil samples from the Upper East region, mean Cd concentrations in soils from the Northern region ranged from 0.014 ± 0.003 mg/kg for the RU fields to 0.042 ± 0.019 mg/kg for the IL fields. The observed differences in mean soil Cd concentration for the three rice production ecosystems in the Northern region were, however, not statistically significant (Table 5).

Table 5 shows that in all the three rice production ecosystems, the Northern region recorded a relatively higher mean Pb concentration than the Upper East region. A *t*-test performed to compare the Pb concentrations between the two regions, however, showed that the differences were not statistically significant (P = 0.10). One-way ANOVA of the soil Pb data also showed no statistical differences between the rice production ecosystems in both the Northern and Upper East regions (Table 5).

### 3.4 Elemental concentrations in rice grain samples

In the Northern region, rice grain As concentration was highest in the RU fields with a mean of 0.005 ± 0.001 mg/kg (Table 6). Rice grains sampled from the Upper East region had higher mean concentrations of As, which ranged from 0.061± 0.002 mg/kg for the RU fields to 0.063 ± 0.003 mg/kg for the RL fields and to 0.064 ± 0.006 mg/kg for the IL fields (Table 6). For both regions, one-way ANOVA showed no significant difference between rice production ecosystems with regard to rice grain As concentration (Table 6). A simple *t*-test, however, found the observed differences between the two regions with respect to rice grain As to be significant (*P* **<** 0.001).

**Table 6.** Elemental concentrations in rice grain from different production ecosystems.

|  | Region | Rice Production system |  |  | *P*-value |
| --- | --- | --- | --- | --- | --- |
|  |  | Rainfed Upland | Rainfed Lowland | Irrigated Lowland |  |
| Grain As<br>(mg/kg) | Northern | 0.005 ± 0.001 | 0.004 ± 0.002 | 0.004 ± 0.000 | 0.580 |
|  | Upper East | 0.061 ± 0.002 | 0.063 ± 0.003 | 0.064 ± 0.006 | 0.699 |
| Grain Cd<br>(mg/kg) | Northern | 0.001 ± 0.001 | 0.011 ± 0.006 | 0.023 ± 0.003 | 0.013 |
|  | Upper East | BD | BD | BD | - |
| Grain Pb<br>(mg/kg) | Northern | 0.083 ± 0.015 | 0.092 ± 0.020 | 0.104 ± 0.041 | 0.671 |
|  | Upper East | 0.024 ± 0.022 | 0.077 ± 0.045 | 0.025 ± 0.022 | 0.141 |

BD: Below detection

Rice grain Cd concentration in all samples from the Upper East region were below detection limit (Table 6). In contrast, detectable levels were found in samples from the Northern region with means reaching a maximum of 0.023 ± 0.003 mg/kg for the IL fields. One-way ANOVA performed on the Northern region rice grain Cd data yielded a *P*-value of 0.013 (Table 6), indicating that the differences between the three production ecosystems with regard to rice grain Cd content are statistically significant.

For both the Northern and Upper East regions, one-way ANOVA of the mean rice grain Pb concentrations recorded in the three production ecosystems showed no significant differences (Table 6). However, a *t*-test performed to compare mean Pb concentrations in rice grains from the Upper East region to those from the Northern region showed that the latter were significantly more enriched with Pb (*P* = 0.002). Mean concentration of Pb in rice grains from the Northern region ranged from 0.083 ± 0.015 mg/kg in the RU fields to 0.104 ± 0.041 mg/kg in the IL fields (Table 6).

## 4. Discussion

Rice is known to grow under a wide soil pH range of 4.2 – 8.5 with the most favourable range falling between 5.5 and 7.0 (Fo et al., 2012). The soil pH values recorded in this study were, therefore, within acceptable limits and similar to the range reported by Opuni et al. (2018) for rice fields in the Volta region of Ghana. The soil EC values also fall within the acceptable range for plant growth reported by Munns and Tester (2008).

The concentrations of As and Pb recorded in water samples from the study sites were less or equal to the WHO’s provisional guideline value of 10 µg/l (0.01 mg/l) for both elements in drinking water (WHO, 2011b). Adomako et al. (2010) also found relatively low concentrations of As (0.67 – 3.23 µg/l) in surface waters used for rice irrigation in southern Ghana, except in water from gold mining impacted rice fields in the Ashanti region. With respect to Cd concentration in the water samples, differences in geology and water-rock interactions could account for the observed differences between the Northern and Upper East regions (Baba and Gündüz, 2017). Usually, Cd concentrations in unpolluted natural waters fall below 0.001 mg/l, but phosphate-based fertilizers constitute a major source of diffuse cadmium pollution (WHO, 2011a) and may also account for the observed concentrations in water samples from the Northern region. This hypothesis would, however, require further investigation since fertilizer was applied in both regions. Besides, information on timing and frequency of application was not obtained for the current study.

Soil As, Cd and Pb concentrations recorded in the current study are all within the permissible limits of 0.1 – 55 mg As/kg (Wenzel, 2013); < 10 mg Pb/kg (Davies, 1995) and < 1 mg Cd/kg (Alloway, 1995) for uncontaminated soils. The observed difference between the Northern and Upper East region soils, with respect to As concentration, could be due to differences in their particle size distribution. Soil texture is known to influence soil As content with clayey soils being generally higher in As than sandy soils (Liu et al., 2005). The higher As content of the Upper East region soils is, therefore, attributable to their higher % clay content (Table 3). Similarly, geogenic factors such as weathering of the underlying parent rocks, may account for the Cd in the Northern region soils; since the recorded concentrations were statistically comparable across the three different rice production systems within the region.

Rice grain As concentration showed a trend similar to that observed in the rice field soils, with values being significantly higher in grains from the Upper East region. This could be attributed to a higher transfer of As to rice grain from the Upper East region soils which were also significantly higher in As than the Northern region soils. The observed rice grain As concentrations were, however, below the maximum permissible limit of 0.2 mg As per kg of polished rice set by the Codex Alimentarius Commission (CAC, 2014). Although rice grain Cd concentrations also fell below the maximum limit of 0.2 mg/kg set for polished rice in the European Union (Commission Regulation, 2006), the significant differences across rice production systems in the Northern region (Table 6) indicate that Cd concentrations in soil and irrigation water contributed to rice grain Cd content. It is worthy of note that although rice grain Pb concentrations were also found to be significantly higher in the Northern region than in the Upper East region, the values fell below the maximum limit of 2 mg per kg of cereals set in the European Union (Commission Regulation, 2006). Thus, with regard to As, Cd and Pb concentrations, the rice grains sampled for the current study are safe for human consumption.

## 5. Conclusion

This paper corroborates findings of the Adomako et al. (2010) study on the PTE status of rice production ecosystems in Ghana. It can be concluded that the country’s rice field soils and surface waters used for rice irrigation are generally low in PTE concentrations. The earlier report of low concentrations of As in rice grains produced in non-gold mining areas of the country (Adomako et al., 2010) has also been confirmed by the current study. The low PTE status of the country’s rice production ecosystems, along with the recorded ranges of soil pH and EC values, indicate their suitability to support the expansion of local rice production to meet the targets set in the National Rice Development Strategy and the PFJ initiative. The call to improve appearance and packaging of locally produced rice grains in order to enhance their competitiveness against imported brands (Adomako et al., 2011; Diako et al. 2011; Ehiakpor et al., 2017) is, therefore, valid and worth heeding. Geogenic factors including soil texture and water-rock interactions may, however, enhance As and Cd enrichment in soils and irrigation waters; which could result in higher rice grain As and Cd content. Regular monitoring of PTE concentrations in the country’s major rice production ecosystems would ensure that the grains produced meet global food safety standards.

## Acknowledgement

The authors would like to thank Dr. Samuel Abebrese and Mr. Alex Yeboah (both of CSIR-SARI) for facilitating access to the rice field study sites.

